# Chromosome-level haplotype-resolved genome assembly provides insights into the highly heterozygous genome of Italian ryegrass (*Lolium multiflorum* Lam.)

**DOI:** 10.1101/2024.10.07.616977

**Authors:** Yutang Chen, Jenny Kiesbauer, Dario Copetti, Daniel Frei, Jürg E. Frey, Christoph Grieder, Roland Kölliker, Bruno Studer

## Abstract

Italian ryegrass (*Lolium multiflorum* Lam.) is an important forage grass, providing a major source of feed for ruminants in temperate regions. Due to its highly heterozygous and repeat-rich genome, high-quality chromosome-level genome assemblies are scarce for Italian ryegrass. Here, we sequenced the genome of a genotype from the Italian ryegrass cultivar ‘Rabiosa’ (hereafter referred to as Rabiosa), and we obtained Oxford Nanopore Technologies long reads (∼60-fold coverage), Illumina short reads (∼85-fold coverage) and high-throughput chromosome conformation capture data (∼60-fold coverage). With Rabiosa as the parent, we constructed an F_1_ population consisting of 305 individuals, which were genotyped by reduced representation sequencing for linkage map construction and quantitative trait loci (QTL) analysis. Using whole-genome sequencing data of Rabiosa and the genetic linkage map, we first generated a chromosome-level unphased haploid assembly (scaffold N50 of 338.75 Mb, total BUSCO score of 94.60%). Then, based on the unphased assembly and a reference-based phasing approach, we generated a chromosome-level haplotype-resolved assembly containing both haplotypes (scaffold N50 of ∼250 Mb and total BUSCO score of ∼90% for each haplotype). Between the two haplotypes of Rabiosa, we observed a highly collinear gene order at chromosome level and a high sequence variation at local level. With a graph-based reference built from the unphased and the haplotype-resolved assemblies of Rabiosa, we conducted a QTL analysis, and two QTL significantly associated with stem rust resistance were detected. The genome assemblies of Rabiosa will serve as invaluable genomic resources to facilitate genomic applications in forage grass research and breeding.

## Introduction

Italian ryegrass (*Lolium multiflorum* Lam.) is one of the most important forage grasses widely grown in temperate regions across the world, providing highly digestible and palatable feed for livestock (Boller *et al*., 2010). It is a diploid, outcrossing species containing seven pairs of chromosomes (2n=2x=14) with a very high level of heterozygosity (Copetti *et al*., 2021). Its genome size is estimated to be around 2.5 Gb (Copetti *et al*., 2021), of which 70% are repetitive sequences (Zwyrtková *et al*., 2020). The relatively large genome size, together with the high degree of repetitiveness and heterozygosity, make genome assembly of this species very challenging. As a result, so far only two draft genome assemblies were reported with one being highly fragmented (Knorst *et al*., 2019) and the other being haplotype-redundant (Copetti *et al*., 2021). Besides the two draft assemblies, only one chromosome-level haploid assembly was recently reported for Italian ryegrass (Brunharo *et al*., 2024). However, this haploid assembly contains only half of the genetic information of the heterozygous diploid genome with a high chance of missing the causal alleles for traits of interest. To advance forage grass breeding and research, haplotype-resolved diploid assemblies containing both haplotypes are urgently needed for Italian ryegrass.

Traditionally, when *de novo* assembling a heterozygous diploid genome, the assembly algorithm randomly selects only one of the two alleles to be present in the output sequence, leading to a mosaic haploid assembly with alleles among loci randomly alternating between haplotypes (Chin *et al*., 2016; Garg *et al*., 2021). With half of the genetic variation missing, a mosaic haploid assembly is never the best representation for a highly heterozygous diploid genome. Ideally, a heterozygous diploid genome should always be represented as a haplotype-resolved assembly where alleles from both homologous chromosomes are not only present but also correctly combined or phased (Koren *et al*., 2018a; Garg *et al*., 2021; Cheng *et al*., 2021; Llamas *et al*., 2021). To generate haplotype-resolved assemblies with long-read sequencing technologies such as Oxford Nanopore Technologies (ONT) or Pacific Biosciences (PacBio), so far, three approaches have been developed and adopted: bin-then-assemble, such as trio-binning (Koren *et al*., 2018b) and reference-based phasing (Porubsky *et al*., 2021; Garg *et al*., 2021); assemble-then-bin, such as ALLHiC (Guk *et al*., 2022), HapHiC (Zeng *et al*., 2024) and phasing with pollen or pedigree sequencing data (Bao *et al*., 2022; Serra Mari *et al*., 2024; Sun *et al*., 2022; Zhou *et al*., 2020); and graph-based phasing, such as Hifiasm (Cheng *et al*., 2021) and Verkko (Rautiainen *et al*., 2023). These methods have generated promising haplotype-resolved assemblies, suggesting that it is now feasible to generate haplotype-resolved assemblies also for highly heterozygous grass species such as Italian ryegrass.

Recently, a graph-based human pangenome reference was published (Liao *et al*., 2023), showing the advantage of overcoming reference bias using a pangenome reference instead of a single reference. In plant sciences, more and more pangenomic studies for grass species such as wheat (*Triticum aestivum*, Walkowiak *et al*., 2020), barley (*Hordeum vulgare*, Jayakodi *et al*., 2020), maize (*Zea mays*, Gui *et al*., 2022), rice (*Oryza sativa*, Shang *et al*., 2022) and various species of millet (*Panicum miliaceum*, Chen *et al*., 2023; *Setaria italica,* He *et al*., 2023; *Pennisetum glaucum*, Yan *et al*., 2023) have also emerged. All these studies illustrate the transition from single-reference-dependent genomics to pangenomic approaches. Given its high level of genetic diversity and its importance as a source of forage, pangenomic resources for Italian ryegrass should be generated. Moreover, to pave the way for future pangenomic studies for *Lolium* species, methods such as pangenome graph construction, haplotype-aware read mapping and variant genotyping with the graph-based reference could be tested with Italian ryegrass.

In this work, we first aimed at generating a haplotype-resolved genome assembly for a highly heterozygous Italian ryegrass genotype using ONT long reads with high-throughput chromosome conformation capture (Hi-C) data and a genetic linkage map. Then, based on the resulting haplotype assemblies, we aimed at understanding the high level of heterozygosity in the genome and addressing how the high heterozygosity might affect genome assembly and phasing. Finally, to explore the potential of current pangenomic methods, we aimed at building a graph-based reference based on the haplotype-resolved assembly and conducting QTL analysis for resistance to stem rust with the graph-based reference.

## Results

### Whole-genome sequencing and reduced representation sequencing of Italian ryegrass

A genotype of the Swiss Italian ryegrass cultivar ‘Rabiosa’ (Agroscope, Zurich, Switzerland), hereafter referred to as Rabiosa, was selected for whole-genome sequencing (WGS), and the following sequencing data were obtained: ONT long reads (total length 153 Gb, corresponding to ∼60-fold coverage of the haploid genome; read N50, 21 kb), Hi-C reads (540 million 150 bp pair-end data, corresponding to ∼60-fold coverage of the haploid genome) and Illumina short reads (818 million 150 bp pair-end reads, corresponding to ∼85-fold coverage of the haploid genome) from a previous study (Copetti *et al*., 2021).

As an additional resource for scaffolding and QTL analysis, an F_1_ population was created by crossing Rabiosa with another Italian ryegrass genotype (from the cultivar ‘Sikem’). The F_1_ population was subject to genotyping-by-sequencing (GBS), and on average, 2.6 million 90 bp single-end GBS reads were obtained for each of the 305 individuals in the F_1_ population.

### Chromosome-level unphased haploid assembly of Rabiosa

A chromosome-level haploid assembly (Rabiosa v1) was constructed based on contigs generated by Flye (Kolmogorov *et al*., 2019) with ONT long reads. The raw Flye assembly was first polished with accurate short reads, resulting in contigs with a total length of 3.39 Gb (Table S1) and a contig N50 of 660.61 kb (Table S1). Next, haplotigs in the polished assembly were removed with a customized pipeline, resulting in a haplotig-purged assembly with its total length reduced to 2.75 Gb (Table 1). Based on the haplotig-purged assembly, subsequent scaffolding with a genetic linkage map containing 26,203 informative single nucleotide polymorphism (SNP) markers (Figure S1) and Hi-C data generated a chromosome-level assembly with a scaffold N50 of 275.65 Mb and a 77.82% anchoring rate (2.14 out of 2.75 Gb were anchored to seven pseudo-chromosomes, Table 1).

**Table 1.**
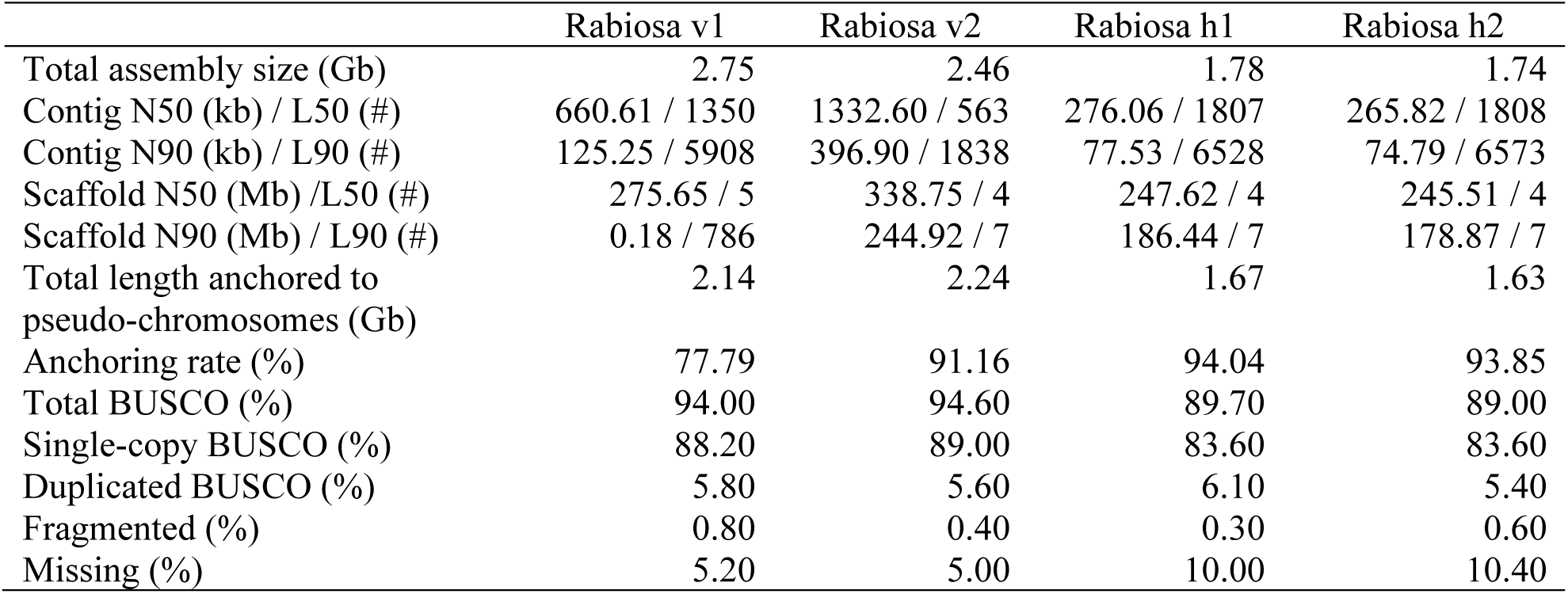
Assembly statistics of Rabiosa assemblies.

In addition to the Rabiosa v1, a second chromosome-level haploid assembly (Rabiosa v2) was generated based on contigs produced by Canu (Koren *et al*., 2017) with the same ONT data. The raw Canu assembly was first polished, resulting in contigs with a total length of 4.18 Gb (Table S1) and a contig N50 of 966.70 kb (Table S1). Haplotigs were then removed from the polished assembly, resulting in contigs with a total length of 2.46 Gb (Table 1) and a contig N50 of 1.33 Mb (Table 1). With the help of Rabiosa v1 as the reference, scaffolding was applied to the haplotig-purged contigs, resulting in a chromosome-level assembly with a scaffold N50 of 338.75 Mb and a 91.16% anchoring rate (2.24 out of 2.46 Gb was anchored to seven pseudo-chromosomes, Table 1).

Both Rabiosa v1 and v2 showed a very high completeness in total BUSCO scores (>=94%, Table 1), and both reached a chromosome-level contiguity. However, as Rabiosa v2 showed a higher contiguity at both contig and scaffold level compared to Rabiosa v1 (Table 1), Rabiosa v2 was chosen as the reference assembly for further analyses in this work.

### Chromosome-level haplotype-resolved assembly of Rabiosa

With the unphased haploid assembly Rabiosa v2, a reference-based phasing approach was applied to separate the reads belonging to the two Rabiosa haplotypes. Seven chromosome-level phase blocks were constructed with high-density SNPs (Figure 1a, Table S2). Based on the haplotypes of these chromosome-level phase blocks, ONT reads were assigned to four groups including haplotype 1, haplotype 2, untagged and unaligned (Figure 1b, Table S3). Roughly 50 Gb ONT long reads (∼20-fold coverage) were assigned to either haplotype, and based on these haplotype-specific read sets, two phased haploid assemblies were generated. Each haplotype assembly showed a total size of ∼1.80 Gb and a contig N50 of ∼270.00 kb (Table 1). By further scaffolding, around 1.60 Gb of each assembly were anchored to seven pseudo-chromosomes, resulting in two chromosome-level assemblies with a scaffold N50 of around 250.00 Mb (Table 1). The two chromosome-level phased assemblies were named Rabiosa h1 and Rabiosa h2, respectively.

**Figure 1.**
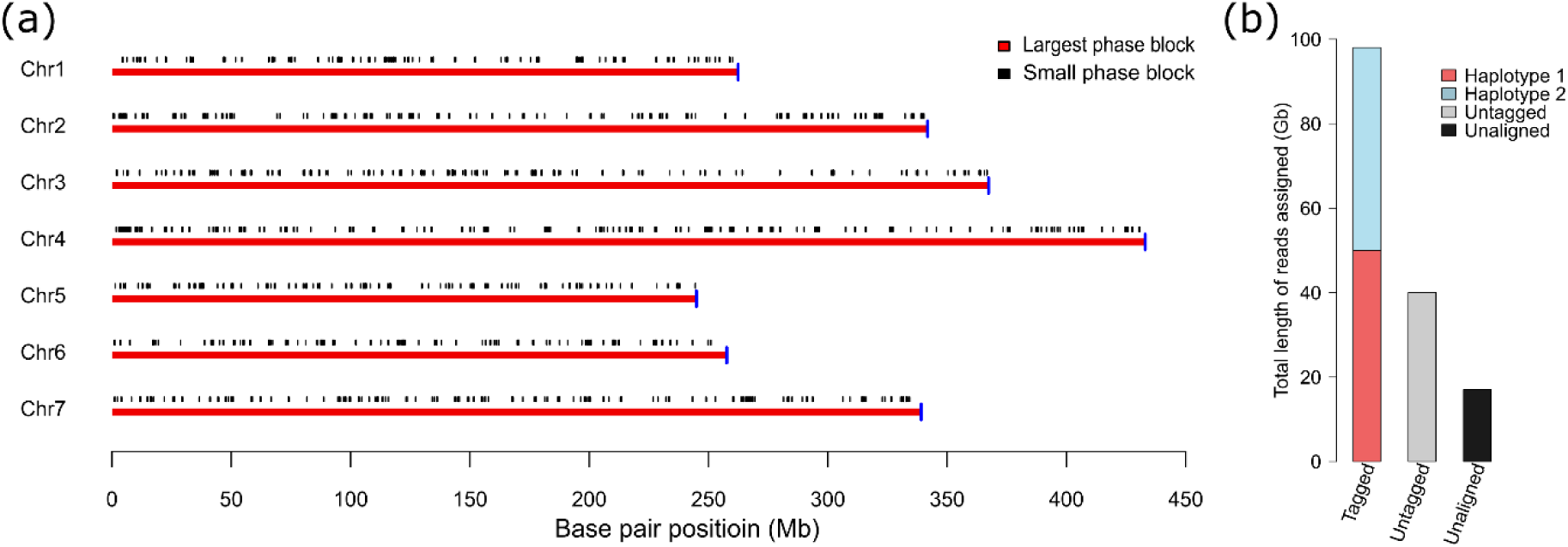
(a) Phase blocks generated by reference-based phasing. The y-axis indicates the seven pseudo-chromosomes (Chr 1-7), and the x-axis indicates the base pair position (in Mb) in the pseudo-chromosome. In each pseudo-chromosome, the red line represents the largest phase block, and black vertical lines indicate small phase blocks that could not be integrated into the largest phase block. The blue vertical line indicates the length of the pseudo-chromosome, showing that the largest phase block spans the whole pseudo-chromosome. (b) Read binning results from reference-based phasing. Reads assigned to haplotypes were classified as tagged, reads mapped to pseudo-chromosomes of Rabiosa v2 but not assigned to either haplotype were classified as untagged, and reads not mapped to any pseudo-chromosomes were classified as unaligned. The sum of tagged, untagged and unaligned reads is equal to the total length of the ONT reads used for reference-based phasing.

### Quality check of the Rabiosa assemblies

Quality assessments were conducted for the unphased haploid assembly (Rabiosa v2) and the two phased assemblies (Rabiosa h1 and h2). All three assemblies showed a high completeness as indicated by their total BUSCO scores (Table 1) and the k-mer comparison results (Figure 2a-c). All three assemblies reached a chromosome-level contiguity as indicated by their scaffold N50 (Table 1). Notably, Rabiosa v2 showed a higher completeness and contiguity compared to the two phased assemblies, reflected by its larger assembly size and higher scaffold N50 (Table 1).

**Figure 2.**
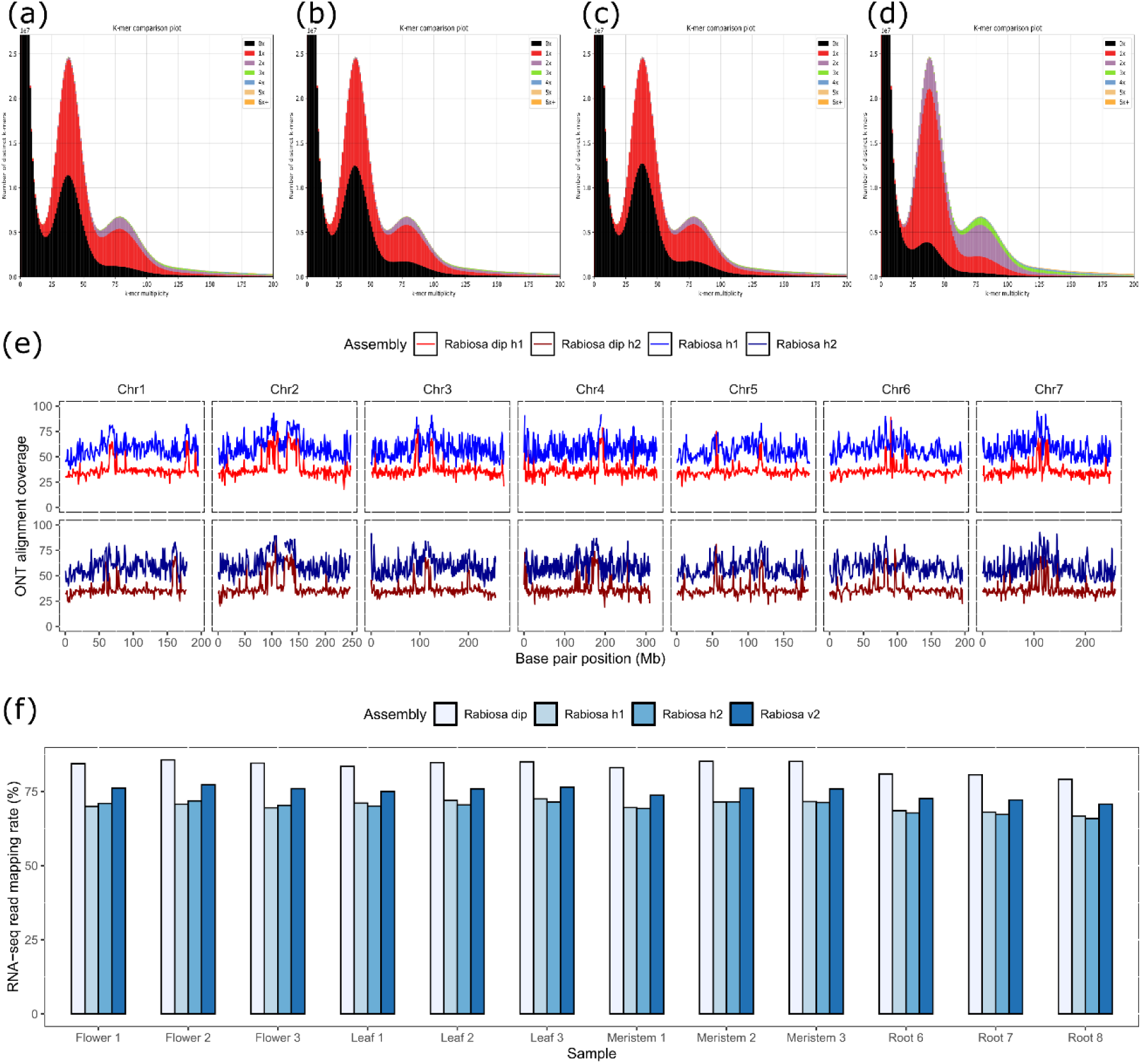
(a)-(d), k-mer comparison between assembly and WGS short reads for Rabiosa v2, Rabiosa h1, Rabiosa h2 and Rabiosa dip, respectively. The plots show whether the presence and copy-number of the k-mers in the assembly match the distribution of the k-mers in WGS short reads. The x-axis represents k-mer frequency in the WGS short reads, and the y-axis shows the corresponding count of the distinct k-mers from the reads. For a heterozygous diploid genome, two peaks are expected for the distribution of the k-mers from reads. The first peak (heterozygous peak) at the lower frequency represents k-mers from the two alleles at heterozygous loci, and the second peak (homozygous peak) at the doubled frequency of the first peak represents k-mers from homozygous loci. The colors in the plot show the copy number of k-mers from the reads in the assembly, with black showing no presence, red showing one copy and purple showing two copies. For a haploid assembly with only one allele present for each locus, the first peak should be half black and half red, and the second should be full red. For a diploid assembly with both alleles present for each locus, the first peak should be full red, and the second peak should be full purple. (a)-(c) suggest that Rabiosa v2, h1 and h2 are haploid assemblies with some homozygous sequences missing (small black area under the homozygous peak). (d) suggests that Rabiosa dip is a diploid assembly with some alleles from heterozygous regions missing (small black area under the heterozygous peak) and some homozygous regions collapsed as one copy (red area under the homozygous peak). (e), ONT read alignment coverage for Rabiosa h1 (blue), haplotype 1 in Rabiosa dip (red), Rabiosa h2 (dark blue) and haplotype 2 in Rabiosa dip (dark red). The read alignment coverage for a haplotype in the diploid assembly is expected to be the half of the alignment coverage of a haploid assembly. (f) RNA-seq read mapping rate (y-axis) in Rabiosa v2, h1, h2 and dip. RNA-seq data were obtained from multiple tissues with three replicates (x-axis).

Rabiosa h1 and h2 were concatenated to form a diploid assembly (Rabiosa dip). Rabiosa dip showed a typical diploid assembly k-mer profile (Figure 2d), suggesting that it was a good representation of the diploid genome. For each haplotype in the diploid assembly, a halved ONT read alignment coverage was observed compared to the haploid assemblies (Figure 2e), suggesting that the haplotypes of Rabiosa were indeed separated. Compared to the three haploid assemblies, Rabiosa dip showed a higher total BUSCO score and, as expected, a much higher duplicated BUSCO score (Table S4) as well as a higher RNA-seq read mapping rate (Figure 2f). This suggested that the haplotype-resolved diploid assembly was a more complete representation of the diploid genome when compared to the haploid assemblies.

The order and orientation of the contigs in the pseudo-chromosomes of Rabiosa v2, h1 and h2 was found to be correct based on the Hi-C contact maps (Figure 3a, b). The structural correctness of the pseudo-chromosomes was also supported by the high consistency between marker order in the genetic map and their physical position in the assembly (Figure 4a, track B).

**Figure 3.**
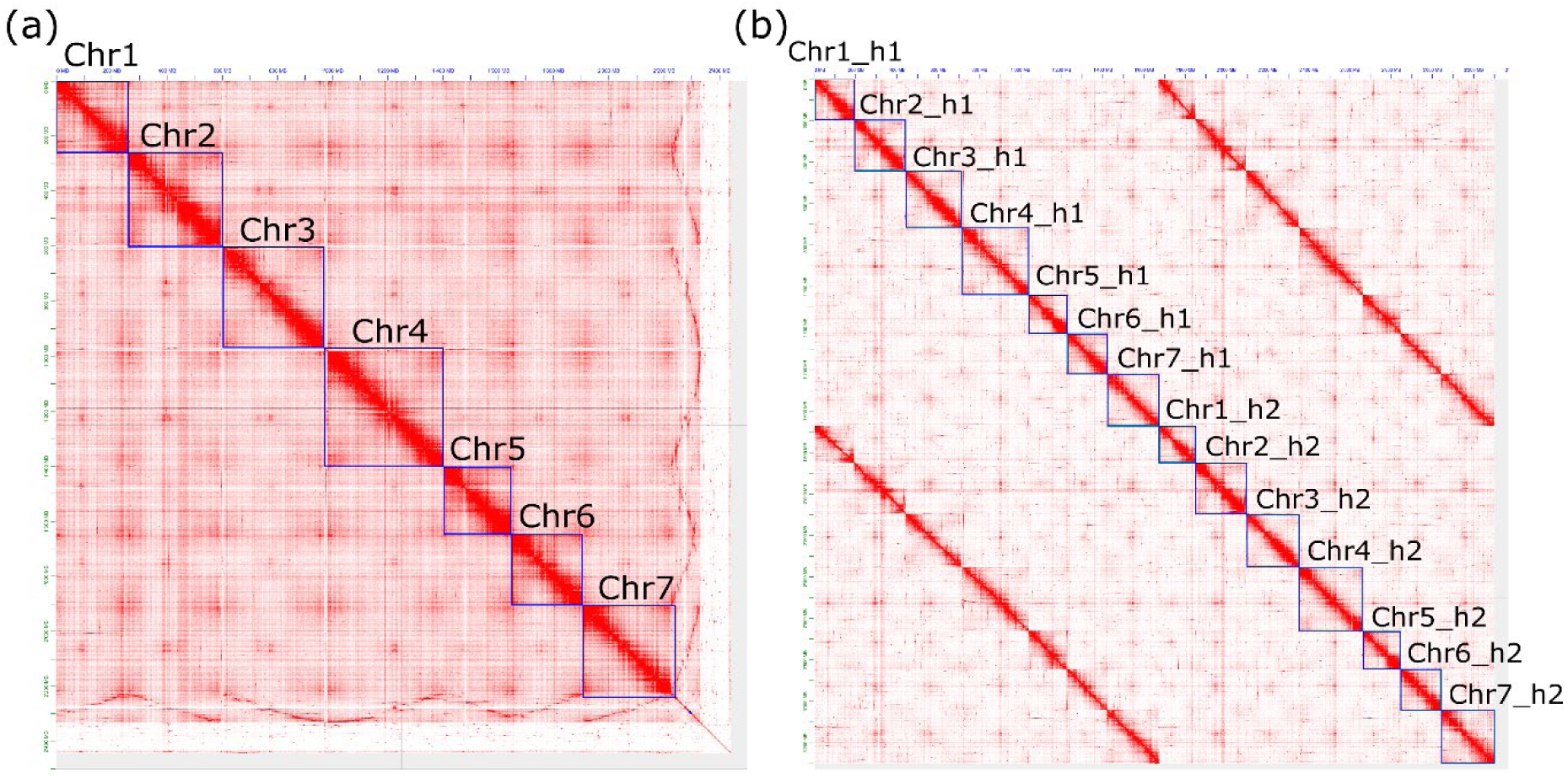
(a) Hi-C contact map of Rabiosa v2. (b) Hi-C contact map of pseudo-chromosomes of both haplotypes in Rabiosa dip. Each red pixel in the plot indicates one interaction between two loci within or between chromosomes, and each blue box represents one pseudo-chromosome. If the sequence in the pseudo-chromosome is correctly ordered and oriented, then more interactions should be observed between closely linked loci within chromosomes, resulting in a “smooth” red diagonal line.

**Figure 4.**
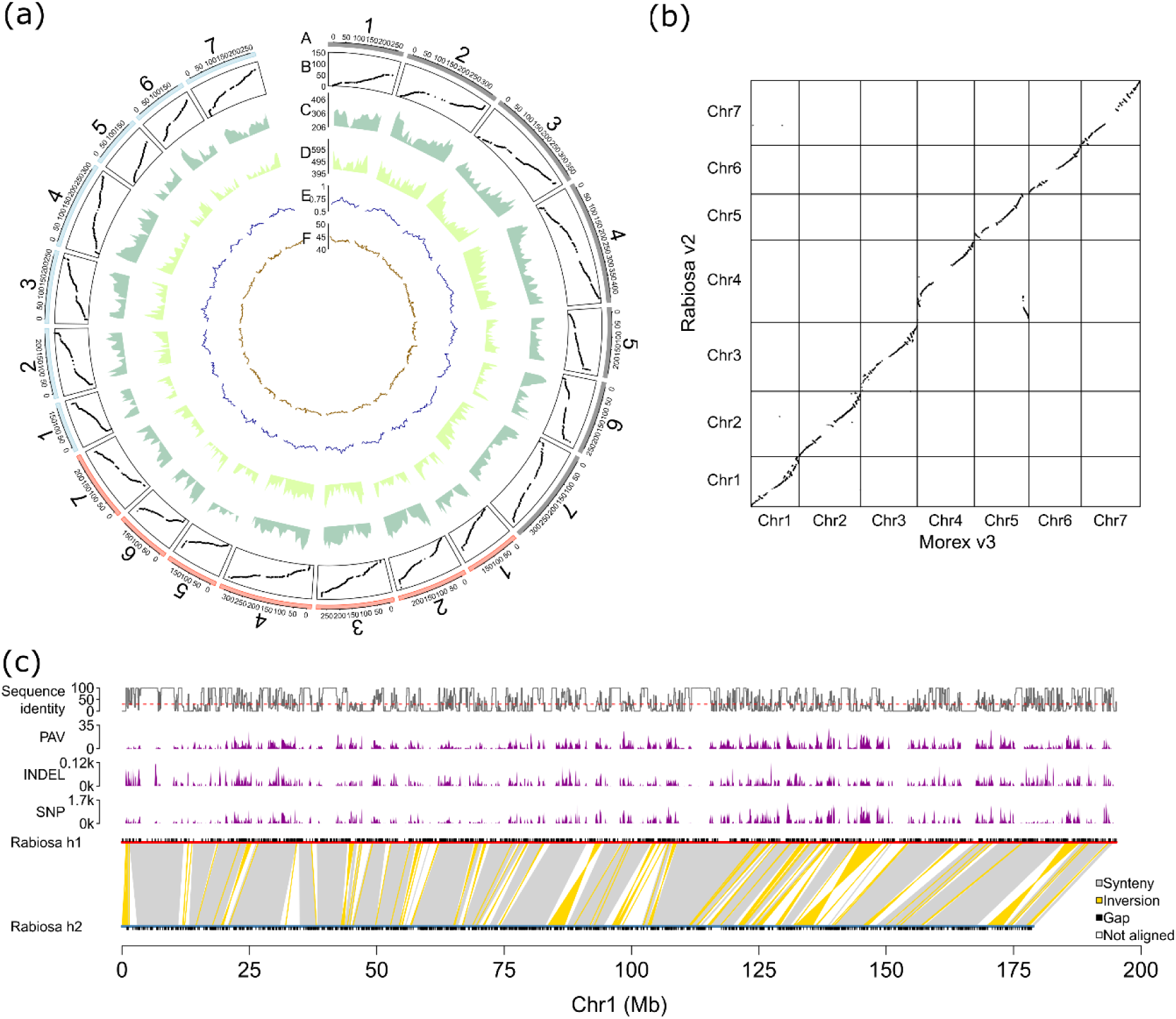
(a) Circos plot showing genomic features of Rabiosa v2, h1 and h2. Track A shows pseudo-chromosomes of Rabiosa v2 (gray), h1 (red) and h2 (blue), respectively, track B the base pair position in the assembly vs genetic linkage map position, track C the number of high confidence genes per 10 Mb, track D the number of low-confidence genes per 10 Mb, track E the fraction of repetitive sequences per 10 Mb and track F the GC content per 10 Mb. (b) genome synteny between Rabiosa and barley cultivar Morex. (c) sequence variation between Rabiosa h1 and Rabiosa h2. The distribution of gaps, SNPs, INDELs, PAVs are displayed. Sequence identity per 100 kb was revealed by the track at the top with the red dashed line indicating 30% identity.

### Genome annotation and gene-based synteny analyses

With an evidence-based genome annotation pipeline, a total of 72,069, 56,142 and 55,928 high-confidence genes were identified for Rabiosa v2, h1 and h2, respectively (Figure 4a, track C). In addition, 113,004, 91,753 and 90,448 low-confidence genes were identified for Rabiosa v2, h1 and h2, respectively (Figure 4a, track D). Around 70% of each assembly was found to be composed of repetitive sequences (Figure 4a, track E, and Table S5), and an average GC content of 44% was observed for each assembly (Figure 4a, track F).

Based on the high-confidence gene models, a highly consistent gene-based synteny was observed between Rabiosa and closely related species including perennial ryegrass (*L. perenne* L.) and barley (Figure 4b, Figure S2). The expected translocation between chromosome 4 of *Lolium* spp. And chromosome 5 of barley (Pfeifer *et al*., 2013) was observed (Figure 4b), validating the correctness of the pseudo-chromosome structure in the Rabiosa assemblies. Notably, the inverted orientation of chromosome 6 and 7 between Rabiosa and Kyuss (Figure S2) should not indicate true chromosome rearrangements between Italian and perennial ryegrass but is indicative for the wrong orientation of pseudo-chromosomes in the first version of Kyuss (Frei *et al*., 2021; Chen *et al*., 2024). A highly consistent chromosome-level gene order was observed between Rabiosa haplotypes (Rabiosa h1 and h2, Figure S3).

### High sequence variation between the two haplotypes of Rabiosa

With a k-mer based analysis, we first estimated the level of heterozygosity of Rabiosa, and a high level of heterozygosity of 3.43% was observed (Figure S5). This suggests a high sequence variation between the two haplotypes of Rabiosa with 3 to 4 SNPs per 100 bp. In parallel to the k-mer based analysis, we also tried to estimate the sequence variation between the two haplotypes of Rabiosa based on not only SNPs but also other types of variants called from whole-genome alignment (WGA) and read mapping. A total of 2,205,246 SNPs, 181,734 small insertions and deletions (indels, length shorter than 50 bp), and 52,578 large presence and absence variations (PAVs, length equal to or greater than 50 bp) were detected between Rabiosa h1 and h2 (Figure 4, Figure Table S6). With these variants, sequence identity between haplotypes was calculated per 100 kb window along the pseudo-chromosomes, and sequence identity as low as 30% was observed between haplotypes in many 100 kb windows (Figure 4c, Figure S4). This low sequence identity suggests that there was a much higher level of sequence variation between the two haplotypes of Rabiosa when structural variants were also considered.

### QTL analysis for stem rust resistance with a graph-based reference of Rabiosa

Based on the three Rabiosa assemblies (Rabiosa v2, h1, h2), a graph-based reference was constructed, which contained 21 paths corresponding to the 21 pseudo-chromosomes of the three assemblies. The total size of the graph-based reference was 3.66 Gb, and there were 243,499,034 nodes and 334,693,667 edges in the graph (Figure 5a).

**Figure 5.**
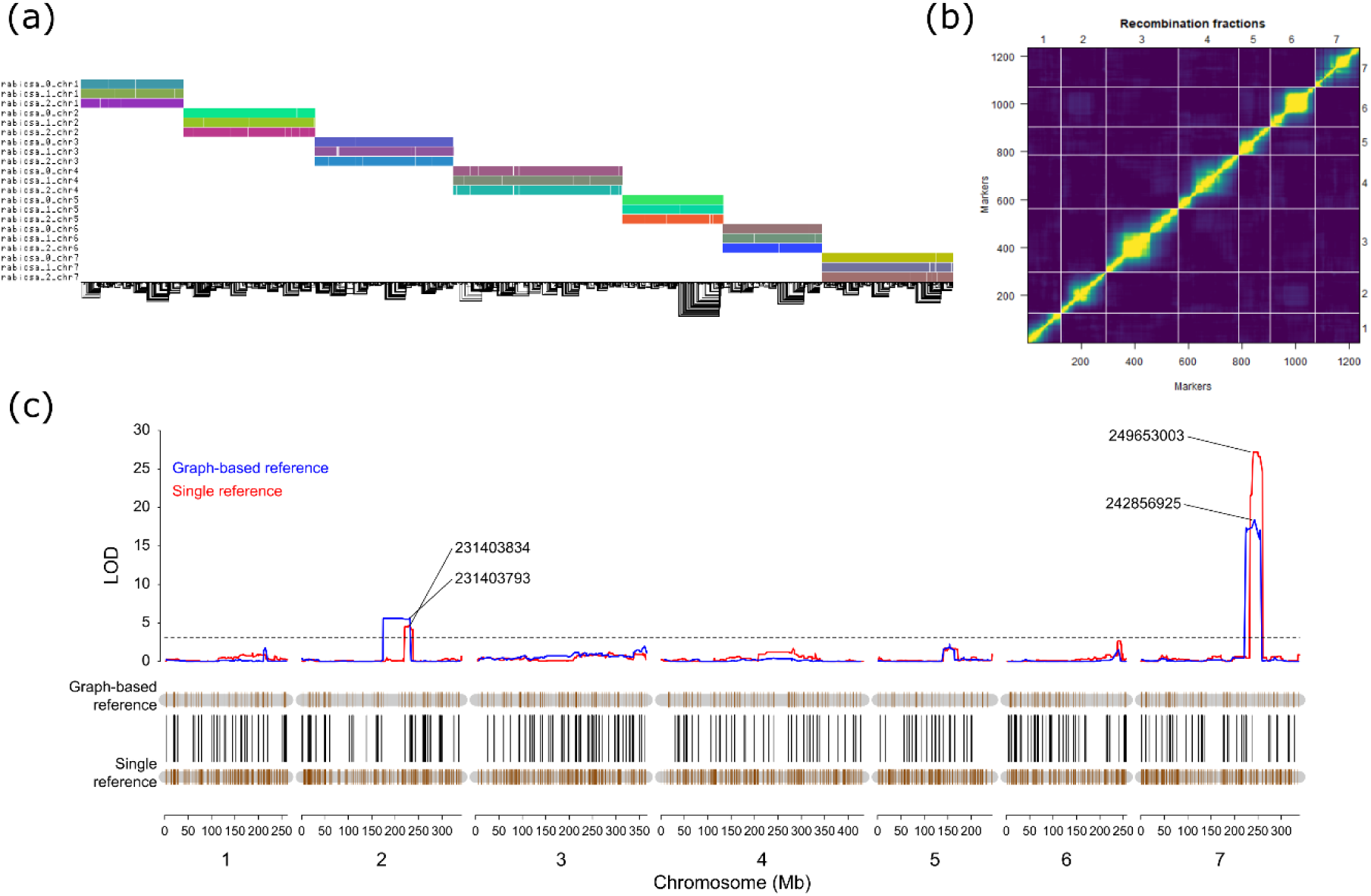
(a) the 1D visualization of the graph-based reference of Rabiosa showing multiple alignment for Rabiosa v2, h1 and h2. Each horizontal colored rectangle corresponds to a pseudo-chromosome in one assembly (one path in the graph) containing nodes in the graph. The name of the pseudo-chromosome was shown by the y-axis label (rabiosa_0, rabiosa_1 and rabiosa_2 correspond to Rabiosa v2, h1 and h2, respectively). The black lines at the bottom are the edges in the graph indicating how nodes were connected. (b) pair-wise recombination fraction between markers in the Rabiosa linkage map generated with SNPs from the graph-based reference. A low recombination fraction (indicated by the yellow color) was observed within linkage groups, and a high recombination fraction (no linkage, indicated by the dark blue color) was observed between linkage groups and distant parts of the same linkage group. (c) results of QTL analysis for stem rust resistance with a graph-based reference and a single reference (Rabiosa v2). The orange vertical lines represent SNPs used for QTL analysis, and black lines connect the same SNP between the graph-based reference and the single reference. The numbers linked to the significant QTL peaks show the base pair position (in Rabiosa v2) of the most significant SNP in the QTL.

Essentially, the graph-based reference is a form of non-redundant multiple sequence alignment, and variants can be directly detected from the multiple sequence alignment. For the graph-based reference of Rabiosa, 54,614 SNPs were obtained based on the position in the unphased haploid assembly (Rabiosa v2). From these SNPs, 1,237 SNPs that were only heterozygous in Rabiosa were selected and used to build a genetic linkage map (Figure 5b). Stem rust resistance was scored for 135 individuals in the F1 population, and a clear segregation of stem rust resistance was observed with a bimodal distribution of the scores (Figure S6). Subsequent QTL analysis for stem rust resistance resulted in two significant QTL, with one located on linkage group (LG) 2 and the other located on LG 7, corresponding to the position in pseudo-chromosome 2 and 7 in Rabiosa v2, respectively (Figure 5c, Table 2). In addition to QTL analysis with the genetic linkage map constructed with the graph-based reference, QTL analysis was also conducted using a genetic linkage map constructed with a single reference (Rabiosa v2, including 5,034 SNPs that were heterozygous in Rabiosa only). The same two QTL that were associated with stem rust resistance were detected (Figure 5c, Table 2), validating the results of the QTL analysis with the graph-based reference. In addition, we also conducted QTL analysis using a linkage map consisting of 6,778 SNPs that were heterozygous in Sikem only. In this analysis, no QTL were detected for stem rust resistance (Figure S7), suggesting that Rabiosa is the source of the stem rust resistance in this mapping population.

**Table 2.** QTL analysis results for stem rust resistance.

| QTL | Reference | Left boundary (bp) | Right boundary (bp) | Highest LOD | LOD threshold |
| --- | --- | --- | --- | --- | --- |
| qtl.chr2 | Rabiosa v2 | 220,462,590 | 237,911,985 | 4.64 | 2.71 |
| qtl.chr7 | Rabiosa v2 | 233,515,340 | 259,843,203 | 27.52 | 2.71 |
| qtl.chr2 | Graph-based | 174,459,966 | 231,403,834 | 5.66 | 2.71 |
| qtl.chr7 | Graph-based | 224,224,230 | 255,175,175 | 18.39 | 2.71 |

## Discussion

Rabiosa v2, the chromosome-level unphased haploid genome assembly of the highly heterozygous Italian ryegrass genotype Rabiosa, showed a much better quality compared to the reported draft assemblies for this genotype (Copetti *et al*., 2021) or other genotypes of this species (Knorst *et al*., 2019). It showed a comparable quality to the recently reported reference-level assembly (Brunharo *et al*., 2024) with regard to the scaffold contiguity and assembly completeness (Rabiosa v2 vs Brunharo’s assembly, Scaffold N50, 338.75 Mb vs 363.00 Mb, total BUSCO score, 94.60% vs 92.80%). Notably, for Rabiosa v2, 60-fold error-prone ONT data were used to generate the assembly, while for the assembly of Brunharo *et al*., nearly 100-fold accurate PacBio HiFi data were used. More importantly, we generated two haplotype assemblies (Rabiosa h1 and h2, Figure 1, Figure 2e, Figure 3b) with the error-prone ONT data, which is more challenging to work with compared to the highly accurate PacBio HiFi data (Li and Durbin, 2024). Although the contigs of each haplotype assembly we generated were rather fragmented (contig N50 270 kb, Table 1), we still managed to anchor more than 93% of the total length of the contigs onto pseudo-chromosomes (Table 1). Besides, when concatenating the two haplotype assemblies into a diploid assembly (Rabiosa dip) and comparing with the haploid assemblies, we found that the diploid assembly was a more complete representation of the gene space of the diploid genome (higher total BUSCO score and RNA-seq read mapping rate, Table S4, Figure 2f). This suggests that a haplotype-resolved assembly should be considered when generating genome assemblies for highly heterozygous species such as Italian ryegrass.

A high level of heterozygosity was estimated with k-mers for Rabiosa (3.43%, Figure S4), which is consistent with the high SNP frequency (1 per 20 bp) observed in ryegrass (Hayes *et al*., 2013). Compared to other published highly heterozygous plant genomes, such as tea (*Camellia sinensis*, heterozygosity 2.31%, Zhang *et al*., 2021), lychee (*Litchi chinensis,* heterozygosity 2.27%, Hu *et al*., 2022) and diploid potato (*Solanum tuberosum*, heterozygosity 2.10%, Zhou *et al*., 2020), Rabiosa showed the highest level of heterozygosity, suggesting that there is a high level of sequence variation between the two haplotypes of Rabiosa. Indeed, a very high level of sequence variation (30% sequence similarity in some 100 kb regions) was observed between the two haplotypes of Rabiosa when estimating the sequence variation with different types of variants including SNPs, indels and PAVs (Figure 4c). Such a high level of sequence variation might explain some of the challenges or technical limitations we encountered when assembling Rabiosa.

We found that the high sequence variation between the two haplotypes of Rabiosa may cause sequence-alignment-based haplotig-purge tools, such as Purge Haplotigs (Roach *et al*., 2018) and purge_dups (Guan *et al*., 2020) to fail to align allelic contigs, leaving more redundant allelic sequences in the assembly after purging (indicated by the duplicated BUSCO score, Table S4). The same situation was observed in other *Lolium* genome assemblies. For example, a haploid *L. rigidum* assembly with 70% duplicated BUSCO score (Paril *et al*., 2022) was generated using Purge Haplotigs (Roach *et al*., 2018), and the new reference-level Italian ryegrass assembly with 20% duplicated BUSCO score (Brunharo *et al*., 2024) was generated using purge_dups (Guan *et al*., 2020). To purge as many haplotigs as possible for Rabiosa, in addition to sequence-alignment-based methods, we also used micro-synteny and BUSCO genes to pair allelic contigs, which led to better purging results compared to using the sequence-alignment-based tools alone (lower duplicated BUSCO scores, Table S4).

Genome phasing was also limited by the high sequence variation between haplotypes. A reference-based phasing approach was chosen in this work to separate the haplotypes of Rabiosa as no parental datasets or highly accurate long reads were available. Based on the unphased haploid assembly (Rabiosa v2) as the reference, we found that around 26% (40 Gb out of the total 153 Gb) of the ONT reads were left unassigned to any haplotypes based on the long-read alignment to the reference. This high proportion of unassigned reads could be attributed to reference bias caused by the high sequence variation between haplotypes rather than sequences originating from homozygous regions (Porubsky *et al*., 2021). As the haploid assembly (Rabiosa v2) was a mosaic representation of the diploid genome with alleles alternating between haplotypes in the pseudo-chromosomes, when mapping long reads from both haplotypes to the reference, reads from one haplotype might be aligned to the reference, but reads from the other haplotype may not be aligned or incorrectly aligned to the reference (Jain *et al*., 2022). If reads were incorrectly aligned to the wrong position due to repetitive sequences (Jain *et al*., 2022), they might be left unassigned to any haplotypes because of no variants called in the repetitive sequences. The high proportion of unassigned reads led to a low coverage (∼20-fold) of haplotype-tagged reads in each haplotype, which compromised the contiguity and completeness of the phased haplotype assemblies (Rabiosa h1 and h2). For the reference-based phasing approach, due to the reference bias caused by the high sequence variation between haplotypes, long reads can only be assigned to haplotypes when they are correctly aligned to the reference. More future work is needed to overcome this limitation when applying reference-based phasing to highly heterozygous genomes. Nonetheless, to the best of our knowledge, the two haplotype assemblies (Rabiosa h1 and h2) generated in this work are the first and the only haplotype-resolved assembly reported for Italian ryegrass so far.

A graph-based reference can be built by adding structural variations (SVs) into a single reference, and this type of graph-based reference has been used for many genome-wide association studies (GWAS) in plants (Liu *et al*., 2020; Zhou *et al*., 2022; Chen *et al*., 2023; Cochetel *et al*., 2023). A graph-based reference can also be built by aligning chromosome-level genome assemblies, and this approach is more accurate than adding a catalog of SVs to a single reference (Hickey *et al*., 2020). However, so far, not many attempts have been made to perform GWAS or QTL analysis in plants with this type of graph-based reference built from *de novo* assemblies (Vaughn *et al*., 2022). To explore the potential of the graph-based reference built from *de novo* assemblies for future pangenomic studies in *Lolium* spp., we generated a graph-based reference with all three Rabiosa assemblies (Rabiosa v2, h1 and h2) and performed QTL analysis for stem rust resistance with the graph-based reference. Based on this graph-based reference, we identified two significant QTL for stem rust resistance, and the same two QTL were also found by the QTL analysis with a single reference (Rabiosa v2, Figure 5c, Table 2). This suggests that the graph-based reference together with the current pangenomic tools, such as the pangenome graph builder (PGGB, Garrison *et al*., 2023), haplotype-aware read mapper (Giraffe, Sirén *et al*., 2021) and the toolkit to manipulate the graph-based reference (vg tookit, Garrison *et al*., 2018) can be reliably applied to highly heterozygous *Lolium* genomes. However, several challenges with the pangenomic approach should be noted. First, due to the high sequence variation between haplotypes, haplotypes are hard to be aligned, and a more suitable whole-genome alignment tool is needed. For example, instead of using the standard aligner wfmash in PGGB, we used AnchorWave (Song *et al*., 2022), which is a synteny-based aligner for pair-wise whole-genome alignment as it is specifically designed for highly heterozygous plant genomes. However, even with AnchorWave, we found that the two haplotypes were not well aligned, leaving the graph-based reference 1.42 Gb larger than the total size of the pseudo-chromosomes of Rabiosa v2 (3.66 vs 2.24 Gb). In contrast, for a pangenome of human built from 90 haplotypes with PGGB, the size of the graph was only 5 Gb, adding just 2 Gb more sequence variations from 90 haplotypes to the haploid genome (genome size 3 Gb, Liao *et al*., 2023). Second, although QTL analysis worked with the graph-based reference, we found that the significant QTL detected on LG 7 with the graph-based reference showed a lower LOD score compared to the same QTL detected with the single reference. This was because fewer SNPs were genotyped in the QTL region with the graph-based reference (Figure 5c). To improve QTL analysis using the graph-based reference, small indels and large structural variations present in the graph-based reference need to be considered in addition to SNPs (Vaughn *et al*., 2022) as SNPs may not be called or genotyped if the two haplotypes could not be aligned.

In conclusion, the chromosome-level unphased haploid assembly (Rabiosa v2) and the two phased haplotype assemblies (Rabiosa h1 and h2) that we generated provide invaluable genomic resources for the forage and turf grass research and breeding community. They will serve as references for Italian ryegrass for genomic applications such as reference-assisted scaffolding, reference-based phasing, QTL analysis and genome-wide association studies. The two QTL linked to stem rust resistance found with the graph-based reference emphasize the potential of graph-based reference and pangenomic tools for future research in forage grasses and other highly heterozygous plant species.

## Methods

### Plant material

A genotype (M.02402/16) of the Swiss Italian ryegrass cultivar ‘Rabiosa’ (Agroscope, Zurich, Switzerland) was selected for whole-genome sequencing in this study. This genotype, referred to as Rabiosa, was crossed with a genotype (M.02002/16) of another Italian ryegrass cultivar ‘Sikem’ (DLF Seeds A/S, Denmark), resulting in a bi-parental population with 305 F_1_ individuals (Methods S1).

### Genome sequencing

ONT sequencing for Rabiosa was conducted following the method described in (Frei *et al*., 2021). Hi-C library preparation and sequencing for Rabiosa was performed at the Functional Genomic Center Zurich (FGCZ), Switzerland. Illumina WGS data of Rabiosa was generated by NRGene as described previously (Copetti *et al*., 2021). For the bi-parental population, GBS libraries were prepared as described by Begheyn *et al*., (2018) and sequenced with Illumina HiSeq 2500 at FGCZ.

### Rabiosa genome assembly

For Rabiosa v1, ONT long reads were first assembled using Flye 2.8.3 (Kolmogorov *et al*., 2019) with --min-overlap 10000 and --iterations 2, then the contigs were polished with accurate short reads using Polca (Zimin and Salzberg, 2020) for one round. After polishing, haplotigs were removed from the polished assembly using a customized haplotig purge pipeline (details see below). Hi-C data were mapped to the haplotig-purged assembly using Arima Hi-C mapping pipeline (https://github.com/ArimaGenomics/mapping_pipeline), and SALSA2 (Ghurye *et al*., 2019) was used to scaffold the haplotig-purged assembly with the Hi-C mapping results. The scaffolds generated by SALSA2 were further anchored to pseudo-chromosomes using ALLMAPS (Tang *et al*., 2015) based on the genetic linkage map generated with the GBS data from the bi-parental F_1_ population (Methods, Genetic linkage map construction). The resulting pseudo-chromosomes from ALLMAPS were used to anchor the scaffolds generated by SALSA2 again using Tritex pipeline (Monat *et al*., 2019) with Hi-C data, producing the Rabiosa v1 assembly.

For Rabiosa v2, ONT reads were first corrected with accurate short reads using FMLRC2 (Mak *et al*., 2023). Then, the corrected ONT long reads were assembled using Canu 2.2 (Koren *et al*., 2017) with batOptions=-eg 0.12 -sb 0.01 -dg 3 -db 3 -dr 1 -ca 500 -cp 50. One round of short-read polishing was applied to the Canu assembly using Polca (Zimin and Salzberg, 2020). After polishing, the same customized haplotig purge pipeline was applied, and then the haplotig-purged contigs were scaffolded to pseudo-chromosomes using Tritex pipeline with Hi-C data and Rabiosa v1 as the reference. For the resulting chromosome-level assembly, a Hi-C contact map was generated using Juicer (Durand, Shamim, *et al*., 2016), 3d-DNA (Dudchenko *et al*., 2017), and manual curation was conducted using Juicebox (Durand, Robinson, *et al*., 2016). This resulted in the Rabiosa v2 assembly.

### GBS data pre-processing

GBS data from the bi-parental F_1_ population was first demultiplexed using Sabre (Najoshi, 2022), and quality check was performed using FastQC (Andrews, 2015). Later, adapters were trimmed off using Trimmomatic (Bolger *et al*., 2014).

### Genetic linkage map construction

Several genetic linkage maps were constructed in this work. The first genetic linkage map constructed was the one used for scaffolding Rabiosa v1. Clean GBS reads obtained from above methods were mapped to the scaffolds generated with SALSA2 (see above) using BWA-MEM (Li, 2013). Then SNPs were called using SAMtools and BCFtools (Danecek *et al*., 2021). Ungenotyped SNPs were imputed using LinkImputeR (Money *et al*., 2017), and the genetic linkage map was built using Lep-MAP3 (Rastas, 2017) by sorting and ordering markers based on the recombination fraction between SNP markers. A second batch of genetic linkage maps were generated based on pseudo-chromosomes in Rabisoa v2 and the haplotype assemblies (Rabiosa h1 and h2, see below), following the same read mapping and variant calling methods. No imputation was done. Lep-MAP3 (Rastas, 2017) was used to construct the genetic linkage maps, and the map distance of the SNP markers was calculated with the given order of the SNP markers from the assemblies. These maps were used to validate the structural correctness of the pseudo-chromosomes in the assemblies and were present in the Circos plot (Figure 4a). For the QTL analysis with the graph-based reference (see below), based on the SNPs obtained from the graph-based reference, a genetic linkage map with SNP markers that were only heterozygous in Rabiosa was constructed using Lep-MAP3 (Rastas, 2017). The position of SNP markers in the genetic linkage map was calculated with the given order of the SNP markers in Rabiosa v2. To verify this genetic linkage map, a recombination fraction plot was generated using R/qtl (Broman *et al*., 2003). For the QTL analysis with the single reference (Rabiosa v2), using Lep-MAP3 (Rastas, 2017), two genetic linkage maps were constructed with SNP markers that were only heterozygous in Rabiosa or Sikem, respectively. Similarly, the position of SNP markers in the genetic linkage maps was calculated with the given order of these SNP markers in Rabiosa v2.

### Customized haplotig purge pipeline

A customized pipeline named PurgeGrass (available at GitHub, https://github.com/Yutang-ETH/PhaseGrass/tree/main/PurgeGrass) was developed to purge both Flye and Canu assemblies in this work. The pipeline pairs allelic contigs based on evidence from all-by-all alignment, micro-synteny and BUSCO genes. All-by-all alignment was conducted using Purge Haplotigs (Roach *et al*., 2018), and micro-synteny between contigs was detected using a pipeline consisting of GMAP (Wu and Watanabe, 2005), GffRead (Pertea and Pertea, 2020), DIAMOND (Buchfink *et al*., 2015) and MCScanX (Wang *et al*., 2012) with transcripts generated in a previous study (Copetti *et al*., 2021). BUSCO genes were detected using BUSCO v4.1.4 (Manni *et al*., 2021) with database embryophyta_odb10. A contig is considered as a haplotig of a longer contig if it fulfills one of the following criteria: 1) having alignment coverage greater than 70% with a longer contig, 2) sharing at least 5 collinear genes with a longer contig, 3) sharing at least one single-copy complete BUSCO gene with a longer contig.

Reference-based phasing and haplotype-resolved genome assembly First, WGS Illumina short reads were mapped to Rabiosa v2 pseudo-chromosomes using BWA-MEM (Li, 2013), and then SNPs were called using SAMtools and BCFtools (Danecek *et al*., 2021). Next, Hi-C data were mapped to Rabiosa v2 pseudo-chromosomes to phase the SNPs using Hapcut2 (Edge *et al*., 2016), following the pipeline described here, https://github.com/vibansal/HapCUT2/tree/master/HiC. This resulted in a sparse chromosome-level phase block per pseudo-chromosome. In parallel, ONT long reads were mapped to the pseudo-chromosomes using Winnowmap2 (Jain *et al*., 2022). By combining the local phase information provided by the ONT read alignment with the long-range phase information provided by the sparse chromosome-level phase block derived from Hi-C data, SNPs were phased through pseudo-chromosomes using Whatshap (Martin *et al*., 2016). This resulted in a dense chromosome-level phase block per pseudo-chromosome. Finally, ONT reads were binned to different haplotypes using Whatshap based on the chromosome-level phase blocks with high-density SNPs, resulting in four sets of reads including haplotype 1, haplotype 2, untagged and unaligned. Only haplotype 1 and 2 reads were used subsequently to generate haplotype assemblies.

Flye (Kolmogorov *et al*., 2019) was used to assemble reads of each haplotype separately. Then one round of short-read polishing was applied to each haplotype assembly using Polca (Zimin and Salzberg, 2020). Next, each assembly was scaffolded with Hi-C using SALSA2 (Ghurye *et al*., 2019) and then mis-joins in the resulting scaffolds were corrected using Tritex (Monat *et al*., 2019). The corrected scaffolds were aligned to Rabiosa v2 to construct pseudo-chromosomes using RagTag (Alonge *et al*., 2022). Additional manual curation to both haplotype assemblies was conducted using Juicer (Durand, Robinson, *et al*., 2016), 3D-DNA (Dudchenko *et al*., 2017) and Juicebox (Durand, Robinson, *et al*., 2016), resulting in Rabiosa h1 and h2. The two haplotype assemblies were concatenated to a diploid assembly (Rabiosa dip), and a diploid Hi-C contact map was also generated with Juicer.

### Assembly quality assessment

The general assembly statistics were calculated with assembly-stats (https://github.com/sanger-pathogens/assembly-stats). The completeness of the assemblies was assessed using BUSCO v4.1.4 (Manni *et al*., 2021) with database embryophyta_odb10. The k-mer profile of each assembly was generated using KAT (Mapleson *et al*., 2017). To calculate the ONT read alignment depth, all ONT long reads were aligned to Rabiosa h1, h2 and dip separately using Winowmap2 (Jain *et al*., 2022), and the alignment depth was calculated using Mosdepth (Pedersen *et al*., 2018). RNA-seq-read mapping was done with the methods described below, and the mapping rate was reported from HISAT2 (Kim *et al*., 2015).

### Evidence-based genome annotation

First, a custom repeat library was generated using RepeatModeler2 (Flynn *et al*., 2020) and ProtExcluder.pl (Campbell *et al*., 2014) based on Rabiosa v2. Then, transposable elements in Rabiosa v2, h1 and h2 were annotated using RepeatMasker (Smit *et al*., 2013) with the custom repeat library.

Gene models were built using EVidenceModeler (EVM) (Haas *et al*., 2008) with evidence from *ab initio* gene prediction, protein alignment and transcript alignment following the method described in Tritex (Monat *et al*., 2019). *Ab initio* gene prediction was performed using BRAKER2 (Brůna *et al*., 2021) with protein sequences from Kyuss (Frei *et al*., 2021). Protein sequences collected from closely related species (Methods S2) were mapped to the assembly using GenomeThreader (Gremme *et al*., 2005), and alignment results were merged using GenomeTools (Gremme *et al*., 2013). Transcripts collected from closely related species (Methods S2) were mapped to the assembly using GMAP (Wu and Watanabe, 2005), and mapped transcripts were merged using Cuffcompare (Trapnell *et al*., 2012) and StringTie merge (Pertea *et al*., 2015). Coding sequences were identified using TransDecoder (Haas, BJ, 2018) with the merged mapped transcripts. Additionally, RNA-seq data of Rabiosa generated previously (Copetti *et al*., 2021) were also used for genome annotation in this work. RNA-seq reads were first trimmed using fastp (Chen *et al*., 2018), then transcripts were constructed using the alignment-based approach with HISAT2 (Kim *et al*., 2015) and StringTie2 (Pertea *et al*., 2015) following the instructions detailed in Pertea *et al*., 2016. All evidence was integrated by EVM to generate consensus gene models.

Using blast (Altschul *et al*., 1990), gene models were searched for against TREP, a transposable elements (TEs) database (Schlagenhauf *et al*., 2016), UniProt reviewed database (The UniProt Consortium *et al*., 2021) and Pfam (Mistry *et al*., 2021) database. Based on the blast results, gene models were classified to high and low confidence groups. High confidence genes were defined as those with a hit in either UniProt or Pfam database and no more than 25% overlap with a TE gene in TREP. In contrast, low confidence genes were defined as those without a blast hit in either UniProt or Pfam database and no more than 25% overlap with a TE gene in TREP. Notably, since the TREP database may not represent all the repetitive sequences in Italian ryegrass, there might be some TE coding genes remaining in the high and low confidence groups. A more comprehensive TE database could be used to improve the classification of the functional protein-coding genes, or TE could be masked prior gene annotation.

Functional annotation was performed using AHRD (https://github.com/groupschoof/AHRD) and InterProScan 5 (Jones *et al*., 2014).

### Gene-based synteny analysis

Using Last (Kiełbasa *et al*., 2011), the high confidence genes of Rabiosa v2, h1 and h2 were aligned against genes from the barley cultivar ‘Morex’ (Mascher *et al*., 2021), Kyuss, a doubled haploid perennial ryegrass genotype (Frei *et al*., 2021), and Lp261, a self-compatible perennial ryegrass genotype (Nagy *et al*., 2022). After alignment, gene-based synteny was constructed using MCscan (Python version; 10.5281/zenodo.594205). The same analysis was applied to check the synteny between Rabiosa v2 and the two haplotype assemblies, and synteny between Rabiosa h1 and h2.

### Calculate sequence identity between two haplotypes

Pseudo-chromosomes of Rabiosa h2 (query) were aligned against pseudo-chromosomes of Rabiosa h1 (reference) using AnchorWave (Song *et al*., 2022) allowing inversions. The resulting alignments were converted to SAM format with maf-convert script provided by AnchorWave. Then the SAM alignment file was further converted to PAF format by paftools.js from Minimap2 (Li, 2018). Next, with the PAF file as input, SyRi (Goel *et al*., 2019) was used to call SNPs, small indels, large PAVs and inversions.

In parallel, short reads were aligned to Rabiosa h1 to call SNPs and small indels with BWA (Li, 2013) and BCFtools (Danecek *et al*., 2021). ONT long reads were aligned to Rabiosa h1 with Winnomap2 (Jain *et al*., 2022), and PAVs were called with Sniffles (Sedlazeck *et al*., 2018). Gaps in Rabiosa h1 and h2 were detected using a custom Python script. Common SNPs, small indels and large PAVs that were detected by both WGA and read alignment were identified by custom R scripts. Common SNPs and indels between two variant calling methods were defined as SNPs and indels sharing the exact start position, while common PAVs between two variant calling methods were defined as PAVs that overlap each other or the start position of one PAV is within ten base pairs downstream the end position of the other PAV. Only common variants were used for sequence identity calculation and PAVs containing gaps were filtered. Sequence identity was calculated per 100 kb along the pseudo-chromosome as (100 kb - total length of common variants - gaps - unaligned sequences reported by SyRi)/100 kb.

### Constructing graph-based reference

Pair-wise whole-genome alignment was first performed among Rabiosa v2, h1 and h2 using AnchorWave (Song *et al*., 2022). Then, the resulting alignments were converted to PAF format as described above, and one PAF file was obtained for each pair-wise alignment. All PAF files were concatenated into one PAF file using a custom R script and then fed into PGGB (the PanGenome Graph Builder, Garrison *et al*., 2023) to construct the graph-based reference.

### Genotyping SNPs in the graph-based reference

SNPs in the graph-based reference were detected by deconstructing the graph-based reference using vg deconstruct (Garrison *et al*., 2018). Subsequently, the resulting vcf file from deconstruction was filtered using vcfbub (https://github.com/pangenome/vcfbub), resulting in a vcf file containing only SNPs. GBS data of each F_1_ individual in the bi-parental population was mapped to the graph-based reference using Giraffe (Sirén *et al*., 2021), and the resulting alignment files were further processed with vg toolkit (Garrison *et al*., 2018). Based on the alignment, any SNPs in the previous vcf file covered by at least five GBS reads of one F_1_ individual were genotyped by vg tookit, resulting in one vcf file with the genotypes of this individual. All individual vcf files were finally merged to one vcf file using BCFtools merge (Danecek *et al*., 2021).

### Phenotyping and QTL analysis

The F_1_ population was grown at the same location (Zurich, Switzerland) each year in a different field for three years from 2018 to 2021 (Methods S1). Resistance to stem rust of 135 F_1_ individual was scored from 1 to 9 according to the severity of the infection of stem rust with 1 being no infection and 9 being the severest infection (Methods S3). The phenotype of stem rust resistance was measured each year at harvesting, and a best linear unbiased estimator (BLUE) was calculated for each genotype with all the phenotypic data collected across years (Methods S3). The BLUEs were used for QTL analysis.

The genetic linkage maps described above and the phenotypic data were used for QTL analysis was performed using R/qtl (Broman *et al*., 2003) with composite interval mapping (CIM).

## Acknowledgements

This study was supported by the European Union’s Horizon 2020 research and innovation program under the Marie Skłodowska-Curie grant agreement No 847585 – RESPONSE. We would like to sincerely thank Ingo Lenk and Elisabeth Veeckman from DLF Seeds A/S for their valuable input and discussion in genome assembly, genome annotation and other bioinformatic analyses. We would also like to thank ISG-HEST at ETH Zurich for providing computational resources as well as their IT service for this work

## Author contributions

Y.T.C. performed all the bioinformatic analyses except the preprocessing of the GBS data and wrote the manuscript. D.C. helped with sequencing data generation, contributed to data analysis and conceptualization and assisted writing the manuscript. D.F. and J.E.F generated and helped analyzing the ONT data. J.K., constructed the GBS library, collected the phenotypic data of stem rust resistance, performed the QTL mapping analysis with the single reference and assisted writing the manuscript.

C.G. created the bi-parental population, helped in phenotyping and data analysis. R.K. completed the preprocessing of the GBS data, supervised the work and assisted writing the manuscript. B.S. acquired the funding, conceived the project, supervised the work and assisted writing the manuscript. All authors revised and approved the final version of the manuscript.

## Data availability

The unphased haploid genome assembly Rabiosa v2 is publicly available on NCBI under the accession: JAUUTY000000000. The two haplotype assemblies are publicly available on NCBI under the accession: JBHEQO000000000. The transcripts used in our haplotig purge pipeline generated previously by *Dario et al.* together with the genome annotation results including the GFF files and the functional annotation results are available at https://doi.org/10.5281/zenodo.13641219. All sequencing data (ONT data, WGS short reads, Hi-C data and GBS data) generated in this work have been submitted to NCBI under BioProject accession: PRJNA990649. The RNA-Seq data generated previously by *Dario et al.* are available on NCBI with the BioProject accession: PRJNA1156394. Code used for bioinformatic analyses in this work can be found at GitHub, https://github.com/Yutang-ETH/Rabiosa.

