## Supplementary Material for "Chromosome-level haplotype-resolved genome assembly provides insights into the highly heterozygous genome of Italian ryegrass (*Lolium multiflorum* Lam.)"

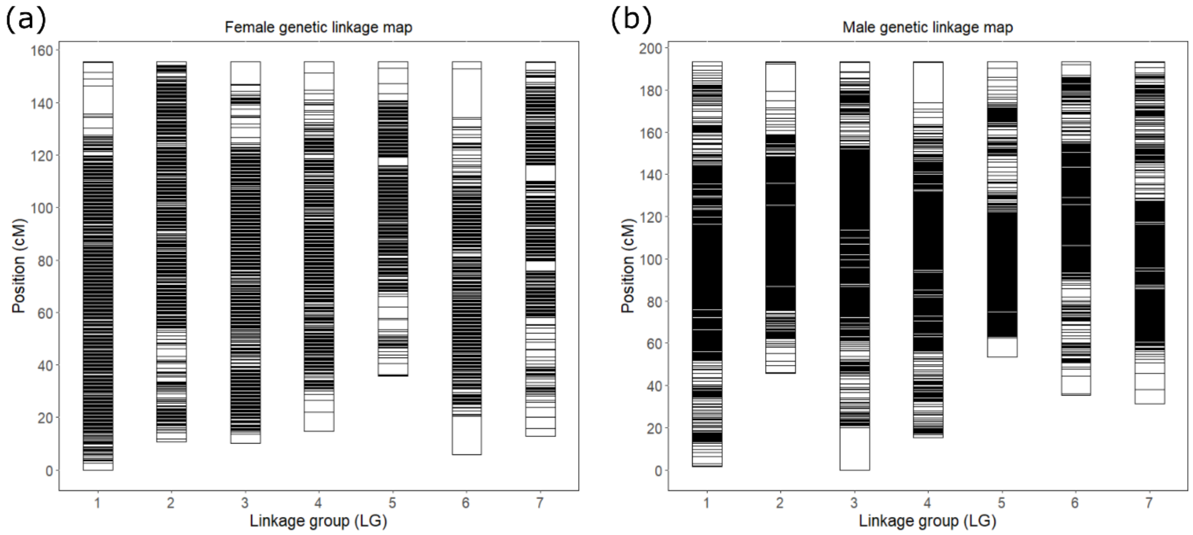


Figure S1. Genetic linkage maps used for scaffolding Rabiosa v1. (a), Female genetic linkage map consisting of 26,203 SNP markers. (b), Male genetic linkage map consisting of 26,203 SNP markers. Each horizontal black segment within the linkage groups represents one SNP marker. Notably, as the F1 individuals were collected from both Rabiosa and Sikem spikes (Methods, S1), the female or the male map does not necessarily refer to the map of Rabiosa or Sikem.

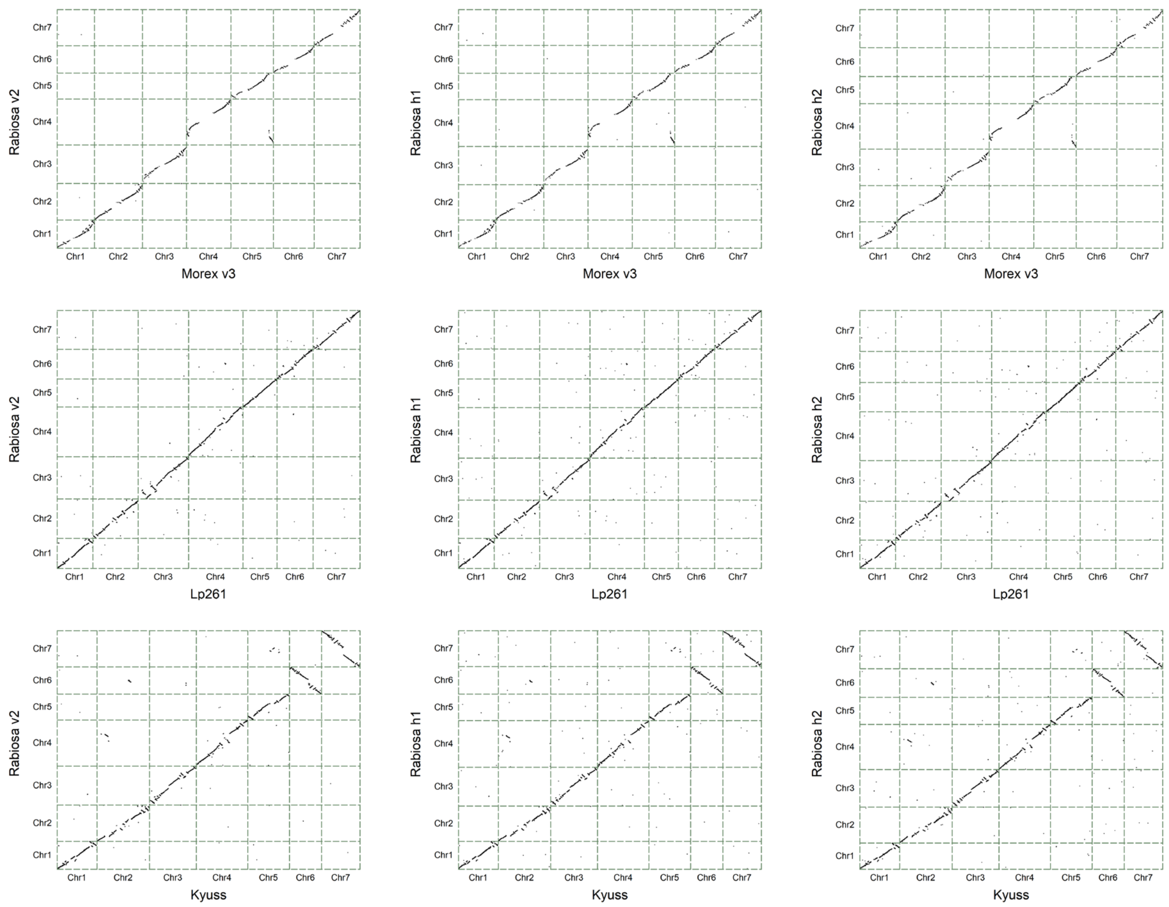


Figure S2. Gene-based synteny between Rabiosa and other genomes. Morex v3 is a barley (*Hordeum vulgare*) cultivar. Lp216 is a perennial ryegrass (*Lolium perenne*) genotype. Kyuss is a doubled haploid genotype of perennial ryegrass (*L. perenne*)

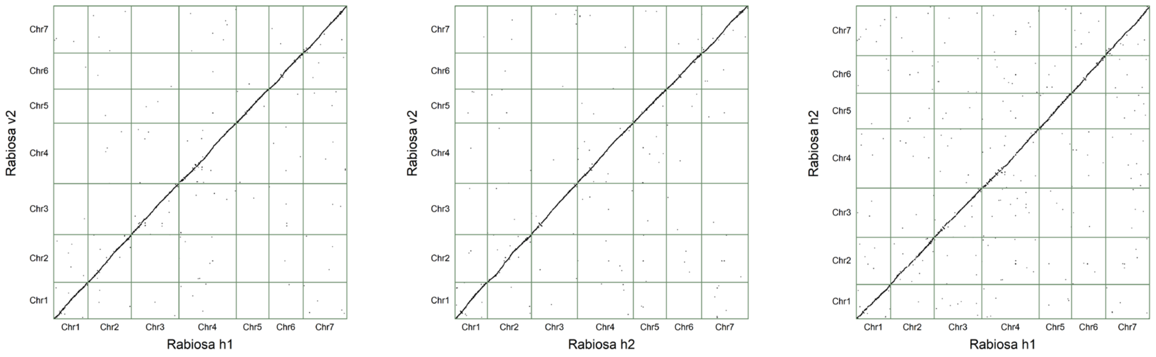


Figure S3. Pair-wise gene-based synteny between Rabiosa v2, Rabiosa h1 and h2

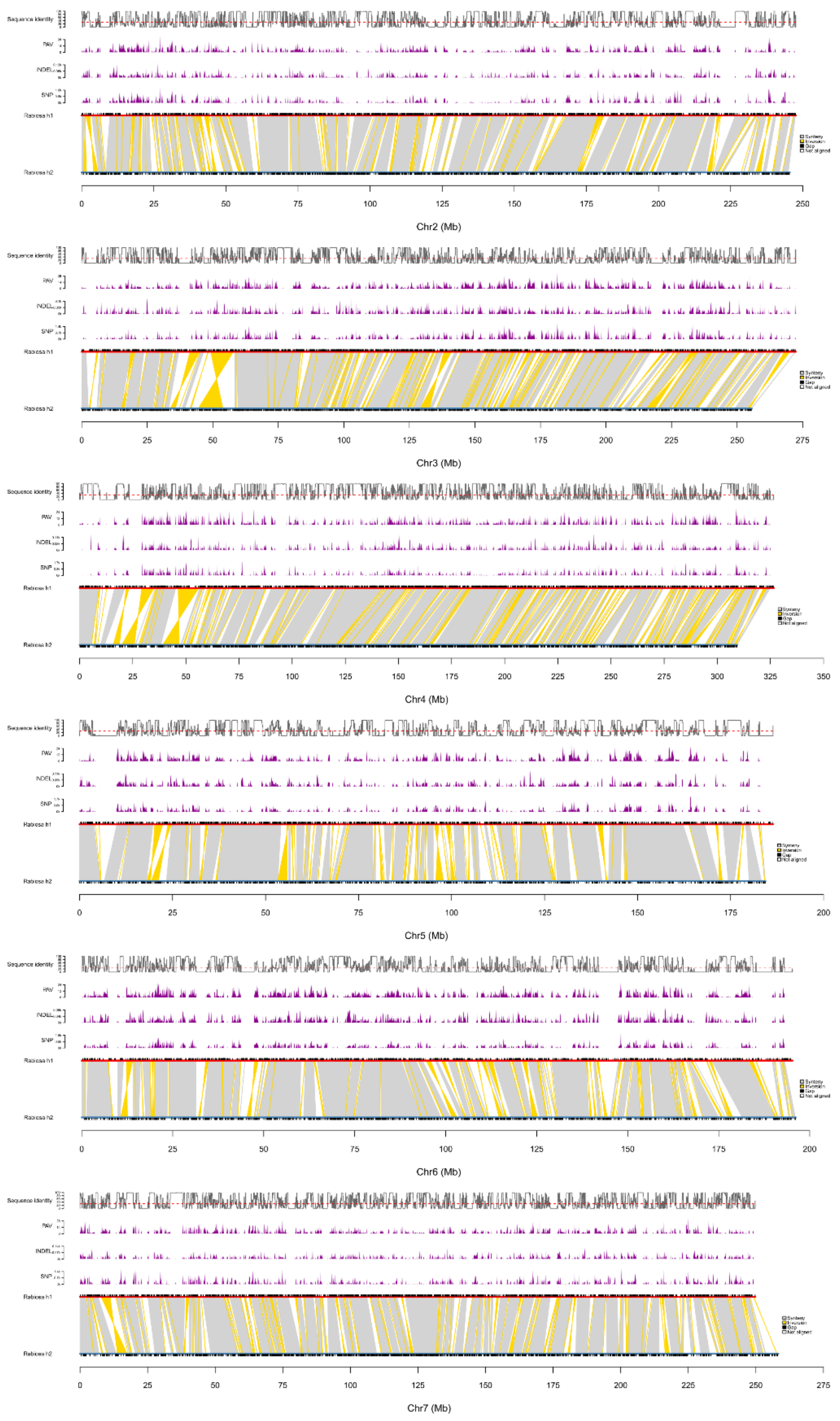


Figure S4. Sequence identity between Rabiosa h1 and Rabiosa h2

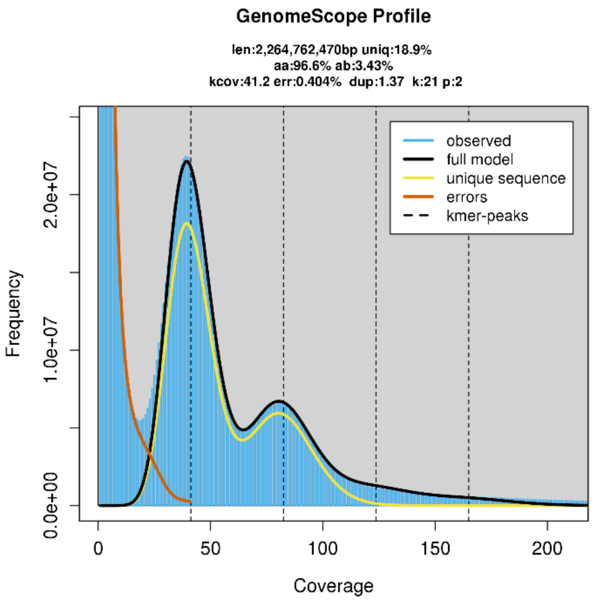


Figure S5. K-mer profile of Rabiosa diploid genome generated using GenomeScope2 with k-mer from whole-genome sequencing short reads.

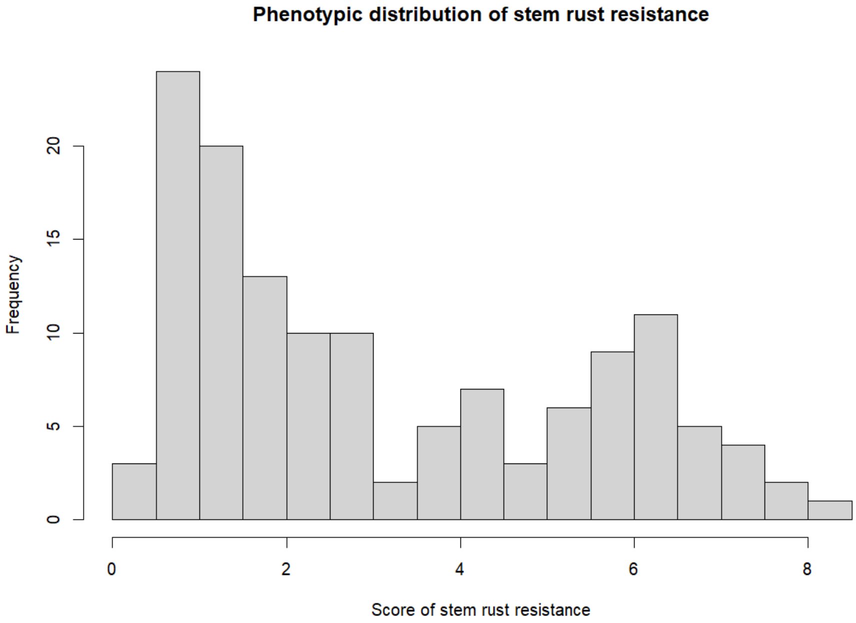


Figure S6. Phenotypic distribution of stem rust resistance based on the score of stem rust resistance from 135 $F_1$ individuals.

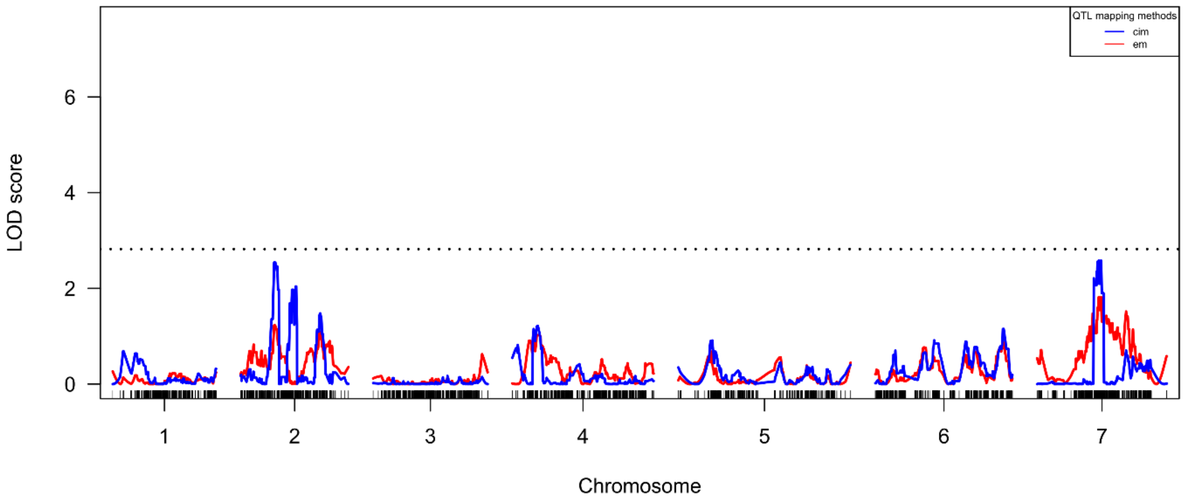


Figure S7. QTL analysis for stem rust resistance with Sikem genetic linkage map. Two statistical methods, including composite interval mapping (CIM) and EM, were used to conduct QTL analysis, and the results from each method were shown in the plot as the blue and the red lines, respectively.

Table S1: Assembly statistics of Rabiosa

| Assemblies | total_length (bp) | number (#) | Gaps (#) | N50 (bp) | L50 (#) | N90 (bp) | L90 (#) |
|---|---|---|---|---|---|---|---|
| rabiosa_h1.fa | 1'779'353'249 | 2'034 | 9'486 | 247'624'434 | 4 | 186'435'459 | 7 |
| rabiosa_h1_polished_contigs.fa | 1'777'003'749 | 11'384 | 0 | 276'062 | 1'807 | 77'529 | 6'528 |
| rabiosa_h2.fa | 1'735'016'293 | 2'062 | 9'479 | 245'508'634 | 4 | 178'866'195 | 7 |
| rabiosa_h2_polished_contigs.fa | 1'732'671'993 | 11'375 | 0 | 265'820 | 1'808 | 74'788 | 6'573 |
| rabiosa_v1.fa | 2'752'220'403 | 7'898 | 5'203 | 275'652'668 | 5 | 178'106 | 786 |
| rabiosa_v1_polished_contigs.fa | 3'392'214'832 | 14'393 | 80 | 660'612 | 1'350 | 125'251 | 5'908 |
| rabiosa_v1_polished_purged_contigs.fa | 2'750'449'603 | 12'783 | 71 | 635'327 | 1'083 | 111'656 | 5'164 |
| rabiosa_v1_polished_purged_salsa_contigs.fa | 2'752'021'603 | 9'886 | 3'215 | 1'935'979 | 351 | 142'075 | 2'547 |
| rabiosa_v2.fa | 2'462'200'656 | 1'918 | 2'299 | 338'745'007 | 4 | 244'921'677 | 7 |
| rabiosa_v2_polished_contigs.fa | 4'179'652'758 | 8'074 | 0 | 966'697 | 1'222 | 226'978 | 4'578 |
| rabiosa_v2_polished_purged_contigs.fa | 2'461'968'756 | 3'383 | 0 | 1'332'597 | 563 | 396'899 | 1'838 |

Table S2: Statistics of Phase blocks

| | Chr1 | Chr2 | Chr3 | Chr4 | Chr5 | Chr6 | Chr7 |
|---|---|---|---|---|---|---|---|
| Total SNPs (#) | 1'064'320 | 1'333'495 | 1'436'084 | 1'693'703 | 1'065'333 | 1'092'690 | 1'359'094 |
| Phased SNPs (#) | 1'056'331 | 1'322'687 | 1'424'888 | 1'680'821 | 1'057'183 | 1'084'431 | 1'347'706 |
| Phased SNPs (%) | 99.25 | 99.19 | 99.22 | 99.24 | 99.23 | 99.24 | 99.16 |
| Unphased SNPs (#) | 7'989 | 10'805 | 11'193 | 12'882 | 8'148 | 8'258 | 11'388 |
| Phase blocks (#) | 776 | 1'042 | 1'010 | 1'380 | 748 | 783 | 919 |
| SNPs in the largest block (#) | 1'050'411 | 1'314'780 | 1'416'897 | 1'668'435 | 1'052'284 | 1'079'145 | 1'341'820 |
| SNPs in the largest block (%) | 98.69 | 98.60 | 98.66 | 98.51 | 98.78 | 98.76 | 98.73 |
| Length of the largest block (bp) | 262'400'071 | 341'736'942 | 367'334'000 | 432'989'167 | 244'874'204 | 257'629'914 | 339'096'701 |
| Length of the pseudo-chromosome (bp) | 264'624'494 | 345'633'772 | 371'026'499 | 442'563'544 | 246'243'529 | 259'232'995 | 341'651'603 |

Table S3: Statistics of binned reads from reference-based phasing

| Bin | ONT reads (#) | Total length of ONT reads (bp) | Average length of ONT reads (bp) |
|---|---|---|---|
| Haplotype 1 | 3'703'084 | 49'511'773'743 | 13'370 |
| Haplotype 2 | 3'571'897 | 47'741'308'009 | 13'366 |
| Untagged | 6'398'652 | 39'766'446'651 | 6'215 |
| Unmapped | 2'003'066 | 16'474'467'909 | 8'225 |

Table S4: BUSCO (v4.1.4) assessment results of genome assemblies

| Assemblies | Complete and single-copy | Complete and duplicated | Fragmented | Missing |
|---|---|---|---|---|
| Flye polished contigs | 26.60% | 72.70% | 0.20% | 0.50% |
| Flye polished contigs purged by Purge Haplotigs | 58.90% | 38.30% | 0.50% | 2.30% |
| Flye polished contigs purged by purge_dups | 58.70% | 36.40% | 0.60% | 4.30% |
| Flye polished contigs purged by PurgeGrass | 85.80% | 8.40% | 0.90% | 4.90% |
| Rabiosa v1 | 88.20% | 5.80% | 0.80% | 5.20% |
| Canu polished contigs | 22.50% | 76.00% | 0.20% | 1.30% |
| Canu polished contigs purged by Purge Haplotigs | 55.00% | 40.80% | 0.40% | 3.80% |
| Canu polished contigs purged by PurgeGrass | 84.10% | 11.00% | 0.20% | 4.70% |
| Rabiosa v2 | 89.00% | 5.60% | 0.40% | 5.00% |
| Rabiosa h1 | 83.60% | 6.10% | 0.30% | 10.00% |
| Rabiosa h2 | 83.60% | 5.40% | 0.60% | 10.40% |
| Rabiosa dip | 23.70% | 72.60% | 0.10% | 3.60% |

Table S5: Transposable Elements (TE) annotation results

| | Rabiosa v2 | | | Rabiosa h1 | | | Rabiosa h2 | | |
|---|---|---|---|---|---|---|---|---|---|
| | Number of elements | Total length (bp) | Percentage of sequence (%) | Number of elements | Total length (bp) | Percentage of sequence (%) | Number of elements | Total length (bp) | Percentage of sequence (%) |
| Class I: Retroelements | 1'004'778 | 1'216'369'770 | 49.40 | 764'884 | 932'095'795 | 52.38 | 742'321 | 906'937'806 | 52.27 |
| LINEs | 75'296 | 41'346'775 | 1.68 | 37'149 | 16'537'021 | 0.93 | 36'610 | 16'120'759 | 0.93 |
| RTE (RIT) | 448 | 103'052 | 0.00 | 358 | 83'940 | 0.00 | 384 | 87'580 | 0.01 |
| L1 (RIL) | 73'349 | 40'705'967 | 1.65 | 35'669 | 15'968'215 | 0.90 | 35'174 | 15'610'630 | 0.90 |
| SINEs | 0 | 0 | 0.00 | 0 | 0 | 0.00 | 0 | 0 | 0.00 |
| LTR elements | 929'482 | 1'175'022'995 | 47.72 | 727'735 | 915'558'774 | 51.45 | 705'711 | 16'120'759 | 51.34 |
| Bel-pao (RLB) | 470 | 40'913 | 0.00 | 419 | 37'659 | 0.00 | 415 | 36'791 | 0.00 |
| Copia (RLC) | 312'617 | 475'784'059 | 19.32 | 246'551 | 363'358'989 | 20.42 | 240'483 | 353'880'354 | 20.40 |
| Gypsy (RLG) | 440'934 | 626'258'793 | 25.43 | 344'952 | 496'007'298 | 27.88 | 331'755 | 482'084'938 | 27.79 |
| Retrovirus (RLR) | 494 | 160'403 | 0.01 | 333 | 103'833 | 0.01 | 393 | 125'742 | 0.01 |
| Class II: DNA transposons | 64'252 | 25'099'582 | 1.02 | 50'404 | 18'551'465 | 1.04 | 50'081 | 18'211'611 | 1.05 |
| hAT (DTA) | 1'767 | 278'837 | 0.01 | 1'537 | 244'367 | 0.01 | 1'489 | 236'803 | 0.01 |
| PIF-Harbinger (DTH) | 9'795 | 2'199'801 | 0.09 | 8'353 | 1'908'413 | 0.11 | 8'455 | 1'929'219 | 0.11 |
| Helitron (DHH) | 403 | 115'975 | 0.00 | 331 | 92'192 | 0.01 | 356 | 100'916 | 0.01 |
| Unclassified | 1'114'996 | 471'026'951 | 19.13 | 703'040 | 236'389'562 | 13.29 | 687'233 | 231'164'744 | 13.32 |

Table S6: Number of variants for calculating sequence variation between the two haplotypes of Rabiosa

| Pseudo-chromosome | SNP (#) | INDEL (#) | PAV (#) |
|---|---|---|---|
| rabiosa h1 chr1 | 251'196 | 20'329 | 6'180 |
| rabiosa h1 chr2 | 311'166 | 25'856 | 7'436 |
| rabiosa h1 chr3 | 387'379 | 29'746 | 9'140 |
| rabiosa h1 chr4 | 408'978 | 32'466 | 9'934 |
| rabiosa h1 chr5 | 218'022 | 18'830 | 5'185 |
| rabiosa h1 chr6 | 302'529 | 23'570 | 6'970 |
| rabiosa h1 chr7 | 326'076 | 30'937 | 7'733 |

Methods S1

Plant material for QTL analysis

A $F_1$ population was created by a reciprocal cross between the genotype (M.02402/16) of the cultivar "Rabiosa" and a genotype (M.2002) of the cultivar "Sikem". In total, 305 $F_1$ seeds collected from both Rabiosa and Sikem spikes were germinated for two days in petri dishes. The seedlings were transplanted in the greenhouse and were grown there for approximately four weeks. Two clonal replicates of a subset of 135 $F_1$ plants were planted in autumn 2018 in the first environment at Zurich (47.427 °N, 8.516 °E) using an alpha design. Because of severe drought issues, only the first replicate of each plant was phenotyped. The detail of phenotyping is described in Methods S3 below. After phenotyping in 2019, the plants were dug out and divided into three clonal replicates and planted in a second environment at Zurich (47.427 °N, 8.516 °E) using an alpha design. After phenotyping, these plants were again transplanted in a third environment at Zurich (47.427 °N, 8.516 °E) using an alpha design and again phenotyped in three clonal replicates in 2021.

Methods S2

The closely related species from which protein sequences were collected for doing genome annotation include the following:

*Brachypodium distachyon*, https://www.ncbi.nlm.nih.gov/datasets/genome/GCF_000005505.3/

*Hordeum vulgare,* https://wheat.pw.usda.gov/GG3/content/morex-v3-files-2021

*L. perenne,* Kyuss
https://datacommons.cyverse.org/browse/iplant/home/shared/commons_repo/curated/Copetti_Kyuss_assembly_annotation_March_2021

*L. perenne,* Lp261,
https://ryegrassgenome.ghpc.au.dk/DOWNLOAD/Lolium_2.6.1/v3_transcripts/PROT/

*Avena sativa,* https://wheat.pw.usda.gov/GG3/graingenes_downloads/oat-ot3098-pepsico

*Seccale cereale,* https://doi.ipk-gatersleben.de/DOI/8afb3971-b5e1-4748-8f0e-1b929ba73248/01369868-8f23-4a21-834f-113b1a9d922d/1/1847940088).

The transcripts used for genome annotation were collected from following sources:
https://www.ncbi.nlm.nih.gov/geo/query/acc.cgi?acc=GSE144460

https://www.ncbi.nlm.nih.gov/geo/query/acc.cgi?acc=GSE78738

https://www.ncbi.nlm.nih.gov/geo/query/acc.cgi?acc=GSE141654

https://datacommons.cyverse.org/browse/iplant/home/shared/commons_repo/curated/Copetti_Kyuss_assembly_annotation_March_2021

https://ryegrassgenome.ghpc.au.dk/

https://zenodo.org/record/832654#.Ynovtp0za70

Methods S3

Phenotyping of stem rust resistance

In spring 2019, the $F_1$ plants were cut after the end of heading. In the second growth, the seed harvesting date was determined based on the cumulative average daily temperature measured 5cm above the ground after the start of flowering for each plant, individually. Once a plant reached the temperature sum of around 590°C, the plant was harvested by cutting with a sickle and phenotyped for stem rust occurrence. Therefore, stem rust occurrence was scored on each plant using a scale from 1 to 9, whereas 1= no rust occurrence, 2 = trace of rust, 3 = 5%, 4 = 10%, 5 = 25%, 6 = 40%, 7 = 60%, 8 = 75% and 9 = more than 75% of the plant is covered with rust (Schubiger and Boller, 2016). To analyze the phenotypic data across environments, the following linear mixed model was used,

$$y_{ijkl} = \mu + g_i + e_j + g_{eij} + b_{lkj} + \varepsilon_{ijkl} \qquad (1)$$

where, $y_{ijkl}$ represents the measurement for stem rust occurrence on a single plant basis, $\mu$ denotes the overall mean, $g_i$ the effect of genotype $i$, $e_j$ the effect of environment $j$, $ge_{ij}$ the interaction effect of genotype $i$ with environment $j$, $b_{lkj}$ the effect of the $l$-th incomplete block nested within the $k$-th complete block and $\varepsilon_{ijkl}$ the residual error. The best linear unbiased estimators (BLUEs) for each genotype across all environments were calculated by using all factors within equation 1 as random, except $b_{lkj}$, which was considered as fixed effect. All the phenotypic data analysis were conducted using R version 4.3.1 (R Core Team, 2023). Package lm4 (Bates *et al.*, 2015) was used for fitting mixed-effect models. Package emmeans (Searle *et al.*, 1980) was used for calculating the BLUEs.
